# Temporal Feasibility Constraints on Wingbeat-Call Synchrony in Actively Echolocating Bats

**DOI:** 10.1101/2025.06.18.660328

**Authors:** Ravi Umadi

**Affiliations:** Lehrstuhl für Zoologie, TUM School of Life Sciences, Technische Universität München, Liesel-Beckmann-Str. 4, 85354 Freising-Weihenstephan, Germany

**Keywords:** Active Sensing, Behavioural Control, Echolocation, Sensorimotor Coordination, Temporal Constraint, Wingbeat–Call Synchrony

## Abstract

Echolocating bats operate within a closed sensorimotor loop in which call emission, echo reception, sensory processing, and motor response are linked by finite propagation delays and bounded response times. Although synchrony between wingbeats and call timing is frequently observed, it remains unclear when such coordination is temporally feasible and when it must necessarily break down. Here, I develop a constraint-based framework formalising how temporal feasibility limits shape wingbeat-call coordination during active echolocation. Building on the responsivity framework, the analysis derives explicit conditions under which call emission remains phase-locked to a cyclic motor rhythm, and identifies regimes in which phase locking becomes progressively infeasible as acoustic delay shrinks and call rate rises during prey approach. Simulations across three motor-control configurations - fixed wingbeat frequency and excursion, dynamically adjusted frequency, and dynamically adjusted frequency and excursion - show that transitions from synchrony to asynchrony arise as necessary consequences of delayed feedback and bounded motor dynamics, rather than discrete changes in behavioural strategy. Increasing motor flexibility extends the synchrony-permissive range of call rates but does not eliminate the feasibility boundary. Simulation ensembles spanning biologically plausible parameter combinations confirm that regime transitions are robust and that asynchronous call phases exhibit structured clustering near the feasibility boundary. Empirical observations of transient decoupling during prey pursuit and the terminal buzz are consistent with these predicted transitions. The results identify temporal feasibility as a governing constraint on echolocation behaviour, clarify how apparent closed-loop coordination can arise without tight motor coupling, and generate testable predictions for when and why wingbeat-call synchrony must fail during prey capture.

## 1 INTRODUCTION

Echolocation in bats is a paradigmatic example of active sensing, in which an animal actively structures its sensory input through self-generated signals and adapts behaviour based on the resulting sensory feedback [1–9]. During flight, echolocating bats must coordinate signal emission, echo reception, neural processing, and motor actions within a tightly coupled sensorimotor loop, often while navigating complex and dynamically changing environments. This coordination enables rapid localisation of prey, obstacle avoidance, and agile manoeuvring, frequently on millisecond time scales [4, 10–13]. More generally, echolocation belongs to a broader class of active sensing systems in which behaviour continuously shapes incoming sensory information while operating under delayed feedback. In engineered control systems and robotics, similar problems arise when actions influence future observations through propagation delays and bounded processing times, requiring controllers to operate within feasibility limits imposed by system dynamics [14–16]. Comparable interactions between rhythmic motor activity and delayed sensory input are also characteristic of biological sensorimotor loops, where perception and action are tightly coupled through ongoing body–world interactions [17, 18]. These parallels suggest that coordination patterns observed in animal behaviour may sometimes reflect fundamental timing constraints inherent to closed-loop sensing systems rather than specific behavioural objectives. Because call production, respiration, and wing motion share biomechanical and physiological pathways, early work proposed that echolocation calls are intrinsically coupled to the wingbeat and respiratory cycles during flight [19, 20]. This view was reinforced by observations of phase-locked call emission during steady flight and by arguments that synchronising vocal output with expiratory phases could reduce energetic cost [20, 21]. Under this interpretation, wingbeat–call synchrony emerges as an energetically efficient solution for biosonar production.

However, subsequent studies across species and behavioural contexts have demonstrated that such synchrony is neither obligatory nor universal. In *Eptesicus fuscus*, calls are often phase-locked to the upstroke during cruising flight but become increasingly decoupled during feeding buzzes [22]. Field recordings further show that wild bats transiently abandon wingbeat–call coupling in cluttered environments or during manoeuvring, where flexible call timing improves echo reception [23]. More recently, Xia et al. [24] demonstrated that while coupling is common at low duty cycles, it systematically degrades as call rates increase, revealing substantial phase flexibility and call insertion outside stereotyped respiratory phases.

Together, these findings indicate that wingbeat–call synchrony is context dependent and tends to break down when sensory demands increase. Notably, this breakdown occurs precisely in behavioural regimes characterised by high call rates, rapid changes in target distance, or dense clutter—conditions under which the temporal demands of the echolocation loop intensify.

Recent analyses have further suggested that bats may regulate call timing according to spatial heuristics. Jakobsen et al. [25] reported that in aerial-hawking vespertilionid bats, call rate scales with approach velocity in a manner consistent with maintaining an approximately constant distance travelled per call. They proposed a “dual convergence” in sonar sampling, preserving both a constant spatial sampling interval and a stable temporal window to traverse the sonar range. Such observations raise an important question: are patterns in call timing governed primarily by spatial objectives, or do they instead reflect more fundamental temporal constraints?

At present, this question remains unresolved. Most analyses describe call timing in relation to distance, velocity, or behavioural phase, but do not explicitly ask whether the required coordination of call emission, echo return, neural processing, and response is temporally feasible within bounded biological response windows. As a result, observed regularities in call timing may sometimes be interpreted as control objectives even when they could arise more simply from permissive timing conditions.

Here I argue that a central, underexplored constraint governing echolocation behaviour is *temporal feasibility*: the requirement that sensory feedback and subsequent motor responses must be ordered and completed within finite temporal margins to sustain closed-loop interaction. Importantly, the presence of a call–echo–call sequence alone does not establish strict closed-loop control, as apparent feedback coupling can arise trivially when acoustic propagation delays are short relative to biological reaction times. In such buffered regimes, calls may be generated according to internal schedules or kinematic heuristics while still receiving informative echoes before the next emission becomes mechanically or neurally possible. This pseudo–closed-loop operation can mimic echo-anchored control without requiring that call timing be causally governed by echo arrival.

As call rates increase and echo streams lengthen, however, this buffer collapses. Call scheduling then becomes explicitly constrained by the timing of sensory feedback itself: a call cannot be meaningfully emitted before sufficient echo information is available via the previous echo, but call production does not need to await full termination of the echo stream. Instead, call production is organised relative to the informative portion of the echo response, creating a finite temporal window within which action updates must occur. Under such conditions, internal pacing policies are no longer sufficient, and call timing becomes rate-limited by echo arrival and processing delays.

The present study formalises this transition using the *responsivity framework* [26], which abstracts echolocation as a sequence of causally ordered events—signal emission, echo return, neural processing, and behavioural response—each associated with characteristic delays. This formulation yields explicit feasibility conditions under which call scheduling can remain phaselocked to cyclic motor rhythms such as the wingbeat, and conditions under which progressive phase slip, that is, the accumulation of phase difference between the call cycle and the wingbeat cycle across successive emissions, and eventual decoupling become unavoidable (Fig. 1). Crucially, these transitions arise from first-principles timing constraints and do not require changes in behavioural strategy, motivation, or energetic optimisation. In this view, the breakdown of wingbeat–call synchrony is not a failure of coordination, but a predictable and adaptive consequence of finite temporal feasibility.

**Figure 1.**
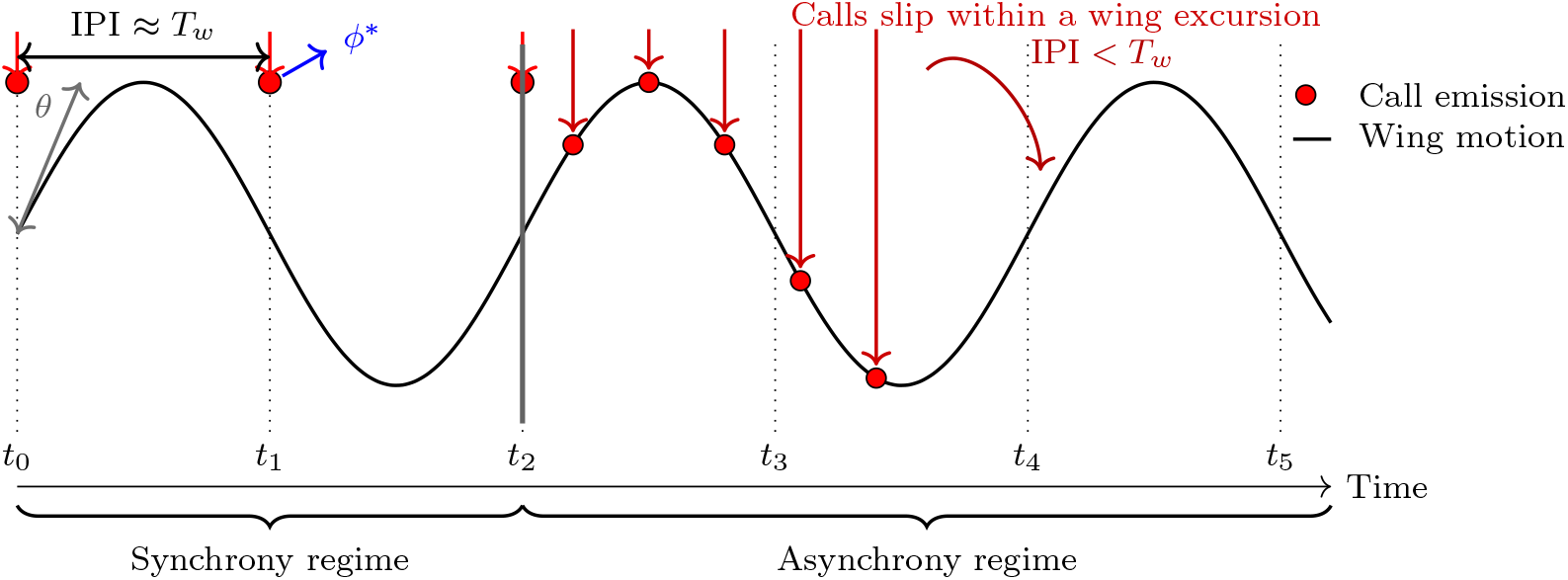
Schematic illustration of the transition from wingbeat–call synchrony to asynchrony in echolocating bats. The sine wave depicts wing motion over time, with red dots marking call emissions. The wingbeat period is denoted by *T*_*w*_, and the interpulse interval between successive calls is denoted by IPI. In the *synchrony regime* (*t*_0_–*t*_2_), calls are phase-synchronised to a consistent point within the wingbeat cycle such that IPI ≈ *T*_*w*_. As echo-processing demands increase, the interpulse interval shortens relative to the wingbeat period (IPI *< T*_*w*_), initiating the *asynchrony regime* (*t*_2_ –*t*_5_) in which successive calls slip in phase and are no longer strictly tied to the wingbeat cycle. The schematic illustrates that synchrony is only possible when the temporal constraints governing call timing remain compatible with the wingbeat rhythm; as sensory demand increases, asynchrony emerges as a necessary consequence of maintaining temporal precision in the echolocation loop.

More broadly, this work situates bat echolocation within a general class of closed-loop active sensing systems subject to delay and feasibility constraints. By treating temporal feasibility as a governing constraint, the present framework provides a unifying account of when synchrony can be maintained, when it must break down, and why such breakdowns can enhance rather than impair sensory performance.

## 2 METHODS

### 2.1 Analytical framework and modelling assumptions

The analytical framework developed here treats echolocation call timing and wingbeat dynamics as two distinct cyclic processes with time-varying frequencies. Call emission is governed by a sensory-driven timing loop whose rhythm is constrained by acoustic propagation delays, neural processing times, and response latencies, as formalised in the responsivity framework [26]. The responsivity coefficient *k*_*r*_ encapsulates the speed at which sensorimotor feedback governs call timing — specifically, it scales the delay between echo reception and the subsequent call emission relative to the concurrent echo delay (Eq. 2). In contrast, wingbeat dynamics arise from biome-chanical and aerodynamic constraints associated with flight, including muscle physiology, body inertia, lift generation, and energetic demands, and are not assumed to be directly controlled by sensory feedback on a call-by-call basis. As a result, the two oscillatory processes are treated as *functionally independent* in their generative mechanisms, even though their interaction gives rise to observable coordination patterns.

This perspective differs from classical coupled-oscillator models, in which synchrony emerges from explicit bidirectional coupling terms or mutual phase adjustment, often with fixed or delayed interactions. Here, no such coupling is assumed. Instead, coordination—or its breakdown— emerges from temporal feasibility constraints imposed by delayed sensory feedback acting on one oscillator (call timing) in the presence of bounded dynamics in the other (wingbeat rhythm). The analysis therefore focuses on when phase-locked coordination is temporally admissible given the ordering and latency structure of the closed-loop system, instead of how oscillators entrain one another. The following sections formalise these constraints and derive the conditions under which synchrony, phase slip, or decoupling must occur.

## 3 THEORETICAL FRAMEWORK

### 3.1 Echo delay and salient reflectors

Let *d*(*t*) denote the instantaneous range to a salient reflector that dominates the relevant echo stream for behavioural guidance, and *c* the speed of sound. In the simplest stationary-reflector case, the two-way acoustic delay *T*_*a*_ is defined as

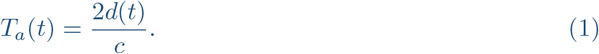

For moving targets or non-negligible bat motion during call emission, higher-order corrections can be included, but the present analysis depends only on the monotonic relation between *T*_*a*_ and the effective range to the echo that the bat uses for feedback. This framing is also consistent with laboratory settings where the farthest prominent reflector within the enclosure can set an upper bound on the longest feasible inter-pulse interval (IPI) – defined as the time between consecutive call emissions – under closed-loop operation.

### 3.2 Responsivity relations and interpulse timing

Within the responsivity framework [26], behavioural coordination is governed by the temporal ordering of four events: call emission, echo return, sensory processing, and initiation of the next action. The present analysis abstracts the latter two stages into a single behavioural response window, denoted *T*_*b*_, which represents the minimal time required for extracting behaviourally relevant information from the echo and implementing the subsequent motor update. Importantly, *T*_*b*_ is not assumed to be fixed, nor is it identified with a specific neural or biomechanical process; rather, it serves as a phenomenological descriptor of the closed-loop response latency.

A core assumption of the responsivity framework is that, over a broad operating range, the effective response window scales with the acoustic round-trip delay *T*_*a*_ according to

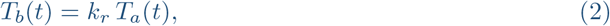

where *k*_*r*_ *>* 0 is a *dimensionless responsivity coefficient*. In th e pr esent st udy, *k*_*r*_ is treated as constant for a given behavioural condition or simulation run: it is not a time-varying control signal, and all temporal variation in *T*_*b*_ arises through changes in *T*_*a*_(*t*). Differences in *k*_*r*_ across simulations or species are interpreted as differences in overall sensorimotor demand or effective response latency, not as rapid within-flight modulation.

At the same time, biological systems are subject to finite processing and actuation limits. Accordingly, the behavioural response window cannot be reduced arbitrarily. I therefore consider the lower bound 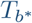, representing the minimum effective response window below which the closed-loop system cannot reliably implement sensory-driven updates. This bound corresponds to the saturation regime of responsivity and reflects irreducible latencies in sensory encoding, decision formation, and motor execution. The response window thus satisfies

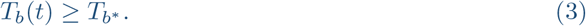

The IPI must accommodate both the acoustic propagation delay and the subsequent behavioural response. Combining Eqs. (2) and (3), the admissible interpulse interval is

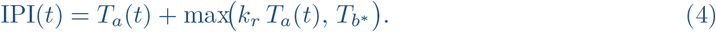

Equation (4) is a *feasibility specification* for closed-loop timing: it defines the shortest interpulse interval compatible with an echo-guided update, given the prevailing acoustic delay and the irreducible response bound.

Two regimes follow immediately. When 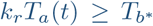, the system operates in a scaling-dominated regime and

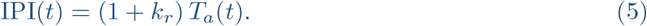

When 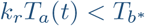, the response window saturates at its lower bound and

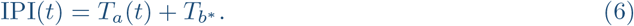

The transition occurs at 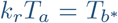 and marks the point at which closed-loop timing becomes explicitly constrained by irreducible response delays rather than by proportional responsivity scaling.

#### 3.2.1 Wingbeat rhythm and phase representation

Let *f*_*w*_(*t*) be wingbeat frequency and *T*_*w*_(*t*) = 1*/f*_*w*_(*t*) the wingbeat period. To discuss coordination in a way that does not assume strict synchrony, it is convenient to represent call times relative to the wingbeat phase. If *t*_*n*_ denotes the time of the *n*-th call emission, the corresponding wingbeat phase can be written as

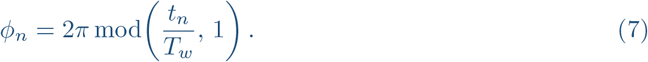

A preference for particular call phases would appear as non-uniformity (clustering) in the distribution of {*ϕ*_*n*_}. In this study, phase is used as a theoretical bookkeeping device to describe feasible coordination regimes, not as a directly inferred quantity.

#### 3.2.2 Feasibility of 1:1 synchrony and higher-order coordination

A common reference point in discussions of motor–vocal coordination is *1:1 locking*, where one call is emitted per wingbeat cycle, implying

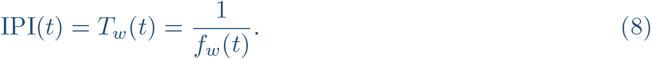

Combining Eq. (8) with the responsivity identity in Eq. (5) yields the wingbeat frequency required for 1:1 locking at a given echo delay,

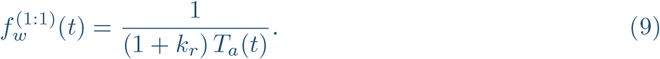

Equation (9) provides a *feasibility condition* for 1:1 locking: if the bat’s achievable wingbeat frequencies do not include 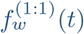 for the relevant *T*_a_ (*t*) and *k*_r_, then strict 1:1 synchrony cannot be sustained unless behavioural adjustments modify the underlying variables entering the feasibility condition. In principle, such adjustments could include changes in wingbeat frequency, flight speed, or call timing strategy that alter the effective relationship between echo delay and call emission. In the absence of such changes, the system must transition to higher-order coordination or phase slip once the required timing falls outside the feasible motor range.

More generally, coordination may occur in higher-order patterns, in which *m* calls occur over *n* wingbeat cycles (an *m*:*n* relation). In such cases,

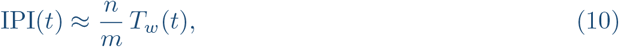

and the corresponding wingbeat frequency compatible with the responsivity-implied IPI is

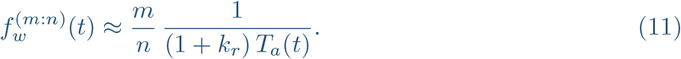

As sensory demand increases (typically as *T*_*a*_ decreases), Eqs. (9)–(11) predict a natural progression from regimes where 1:1 locking is feasible (IPI ≳ *T*_*w*_) to regimes where multiple calls must occur within a wingbeat (IPI *< T*_*w*_), necessitating higher-order locking or phase slip. Importantly, this transition follows from timing feasibility and does not require assuming that bats seek to maintain strict wingbeat synchrony as a behavioural goal.

#### 3.2.3 Coordination regimes implied by temporal feasibility

The relations above imply three broad regimes of behavioural coordination:

1. **Permissive synchrony** (IPI ≳ *T*_*w*_). Temporal feasibility permits 1:1 locking, and calls *may* cluster at a preferred wingbeat phase, consistent with stable coordination under low sensory demand.
2. **Multi-call coordination** (IPI *< T*_*w*_). Multiple emissions occur within a wingbeat cycle, making strict 1:1 locking impossible. Coordination may persist as higher-order locking (*m*:*n* relations) or as phase clustering at multiple phase families — recurring clusters of call-emission phase values within the wingbeat cycle, representing stable locking ratios (e.g., calls always emitted near peak upstroke).
3. **Phase slip /decoupling**. As temporal demand increases further, the higher-order coordinated states described above cease to remain stable. Call-emission phase *ϕ*_*n*_ then drifts systematically across successive wingbeat cycles, producing progressive phase slip and effectively decoupled timing. Consequently, the discrete phase families characteristic of regime 2 dissolve into a broadened or near-uniform phase distribution.

In the remainder of the study, simulations are used to illustrate how transitions between these regimes arise from the responsivity timing relations under changing *T*_*a*_ and bounded wing-beat frequencies. Empirical observations are discussed only insofar as they provide qualitative examples consistent with the predicted regimes.

#### 3.2.4 Phase representation and feasibility of phase-locked call insertion

The following section formalises wingbeat–call coordination in terms of phase compatibility rather than explicit oscillator coupling. To characterise wingbeat–call coordination without conflating oscillatory phase with kinematic amplitude, I represent the wingbeat as a continuous phase oscillator with time-varying frequency. The wingbeat phase is defined as

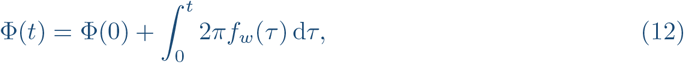

where *f*_*w*_(*t*) is the instantaneous wingbeat frequency. The within-cycle phase is

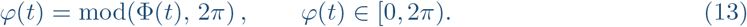

This definition ensures that the phase advances uniformly by 2*π* per wingbeat cycle, independent of wing excursion amplitude.

Wingbeat kinematics constrain *when* within a cycle a call can be emitted in a phase-locked manner. Rather than encoding this constraint through the phase rate, I represent it as an admissible phase subset

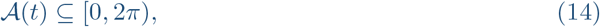

which specifies the range of wingbeat phases at which synchronous call emission is biomechanically permissible. The width and location of *A*(*t*) may depend on wing excursion, flight mode, or other kinematic factors, but its role is purely restrictive: it does not influence the evolution of the wingbeat phase itself.

The responsivity framework yields a feasibility-determined time at which the next call can be emitted based on echo delay and bounded response latency. Denoting this time by 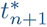, the corresponding predicted wingbeat phase is

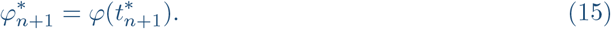

Phase-locked call insertion is feasible if and only if this predicted phase lies within the admissible set:

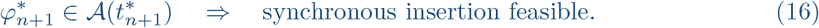

If this condition is violated, no phase-locked solution exists for the given timing constraints, and the call must be emitted asynchronously with respect to the wingbeat:

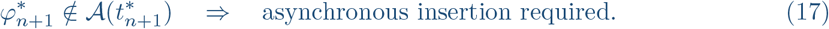

This formulation makes explicit that synchrony and asynchrony are determined by the compatibility between feasibility-imposed call timing and kinematic phase constraints, rather than by an assumed coupling mechanism. Phase slip arises naturally when the admissible phase interval collapses below the timing precision required by closed-loop updating.

#### 3.2.5 Model implications and conditional predictions

The constraint-based framework developed here yields several *conditional implications* for how echolocation call timing may interact with cyclic motor rhythms such as the wingbeat. These implications follow directly from temporal feasibility considerations and do not assume that bats actively seek or maintain synchrony.

1. **Loss of feasible synchrony with decreasing echo delay:** As a bat approaches a target, the acoustic round-trip delay *T*_*a*_(*t*) decreases. Within the present framework, maintaining phase-locked coordination between call emission and a cyclic motor rhythm requires that the period of that rhythm remain compatible with the minimum feasible interpulse interval. When *T*_*a*_(*t*) becomes sufficiently small, this compatibility condition can no longer be satisfied for a given wingbeat frequency. Under such conditions, phase-locked call–wingbeat coordination becomes temporally infeasible, and call timing must transition to a regime that is asynchronous with respect to the wingbeat cycle. This implication is consistent with observations of progressive call–wingbeat decoupling during high call-rate phases, such as the terminal approach to prey [23, 24], without requiring that such decoupling be an explicit behavioural goal.
2. **Context-dependent contraction of the synchrony-permissive regime:** The range of echo delays over which synchrony remains feasible depends on the achievable characteristics of the underlying motor rhythm. If flight conditions or behavioural demands constrain wingbeat frequency or amplitude—such as during rapid manoeuvres, load-bearing flight, or other kinematically demanding tasks—the region of parameter space in which synchrony is possible correspondingly contracts. The model therefore implies that call–wingbeat asynchrony should arise more readily in situations where locomotor degrees of freedom are limited, even if call timing alone would permit synchronous coordination under less constrained conditions.
3. **Morphological and biomechanical modulation of feasibility bounds:** Differences in flight morphology and biomechanics can shift the boundaries of temporal feasibility without eliminating them. Species or individuals capable of sustaining higher wingbeat frequencies or broader kinematic ranges may retain synchrony across a wider range of echo delays and call rates. However, within the present framework, no morphological configuration abolishes the existence of a feasibility threshold: as sensory–motor timing demands increase, a point is eventually reached at which phase-locked coordination is no longer compatible with closed-loop operation.

Taken together, these implications link temporal precision in echolocation to general constraints imposed by delayed sensory feedback and bounded motor rhythms. They suggest concrete avenues for empirical testing, including combined acoustic and kinematic recordings that quantify the evolution of call–wingbeat phase relationships across behavioural contexts, as well as comparative analyses examining how differences in flight mechanics shift the boundaries of synchrony-permissive regimes. Importantly, the framework does not predict specific coordination strategies, but instead delineates the conditions under which particular patterns of coordination are temporally feasible or infeasible.

### 3.3 Wingbeat–Call Synchrony Simulation

To systematically explore the capacity and constraints for maintaining synchrony between wing-beats and call emission, I implemented three distinct simulation regimes:

1. *Constant wingbeat frequency (f*_*w*_): The bat’s wingbeat frequency was held constant throughout the simulated approach, set to a nominal cruising frequency. Call emission timing was determined solely by the echo-processing constraint, independent of the wingbeat cycle. Synchrony was deemed possible only when the calculated call insertion phase (*ϕ*^*^(*t*)) fell within the biomechanical limits of the fixed wingbeat excursion (*θ*), and when no more than one call was required per wingbeat cycle.
2. *Dynamic wingbeat frequency*: Here, the bat’s wingbeat frequency was allowed to increase in step with the required call rate, up to a physiologically defined maximum (*f*_*w*,max_). Synchrony was maintained by locking call emission to the wingbeat cycle as long as *f*_*w*_ ≤ *f*_*w*,max_ and the phase constraint (*ϕ*^*^(*t*) ≤ *θ*) was met. Once the required call rate exceeded this maximum, synchrony necessarily broke down and calls were emitted asynchronously relative to the wingbeat.
3. *Dynamic frequency with adaptive wing excursion (θ)*: Recognising that bats can also modulate the angular excursion of each wingbeat to further increase wingbeat frequency, I implemented a third regime where, upon reaching *f*_*w*,max_, the model allowed the wing excursion angle (*θ*) to decrease dynamically in proportion to the required increase in frequency, down to a set physiological minimum (*θ*_min_), implying an aerodynamic threshold. This adjustment enabled the simulated bat to maintain synchrony further into the high call rate regime. Synchrony, in this context, was defined as the condition in which (i) a single call could be accommodated per wingbeat, and (ii) the predicted call insertion phase fell within the instantaneous wing excursion limit.

Recent field measurements support the assumption that wingbeat kinematics are not fixed during prey pursuit. Using on-board inertial sensors, Stidsholt et al. (2021 & 2025) [23, 27] report systematic variation in both wingbeat frequency and wingbeat acceleration amplitude during natural hunting sequences. Although the IMU-derived amplitude reflects wingbeat intensity rather than excursion angle per se, changes in acceleration and frequency necessarily imply modulation of underlying wing kinematics under finite physiological limits. The motor-control conditions explored here therefore bracket empirically observed regimes: from fixed-frequency motion (Condition 1), through frequency-adaptive wingbeats (Condition 2), to combined frequency and excursion-bound adaptation (Condition 3), without assuming a direct mapping between measured acceleration and geometric excursion.

No separate condition with fixed wingbeat frequency and adaptive wing excursion was included. In the present framework, wing excursion (*θ*) defines the admissible phase window for synchronous call insertion, whereas wingbeat frequency determines the temporal capacity of the motor rhythm itself. If *f*_*w*_ is held constant, varying *θ* alone cannot shorten the wingbeat period or prevent the onset of multi-call-per-wingbeat regimes once the feasible interpulse interval falls below *T*_*w*_. Adaptive excursion therefore does not constitute an independent synchrony-preserving mechanism in this model, but only becomes relevant as an auxiliary degree of freedom when wingbeat frequency is already allowed to increase and subsequently reaches its physiological limit.

Across all regimes, *synchrony* is thus operationally defined as the temporal and biomechanical alignment of call emission with the wingbeat cycle: a call is synchronous if it occurs within a new wingbeat, at a phase within the instantaneous excursion angle (*θ*), and at no more than one call per wingbeat. This framework allows explicit identification of when synchrony is lost—either due to echo-processing constraints, biomechanical limitations, or both—and thus enables predictive testing of the hypothesis that asynchrony becomes inevitable as sensory demands intensify.

#### 3.3.1 Simulation framework and model lineage

The simulations presented here build directly on the *Vespertillio numeralis* implementation introduced in the responsivity framework [26, 28]. In that work, call timing dynamics were governed by echo-delay–dependent responsivity constraints without explicit representation of cyclic flight mechanics. Here, this baseline implementation is extended by incorporating an explicit wing-beat oscillator with bounded frequency and excursion, as described in the preceding sections.

For a detailed description of the original *V. numeralis* model, including acoustic timing, target interaction, and responsivity parameterisation, see Umadi & Firzlaff (2025) [26].

#### 3.3.2 Simulation Parameters

Each simulation was fully specified by a MATLAB parameter structure, params, whose key fields are described in Table 1. All simulation outputs (including synchrony flags, wingbeat frequency, call phase, and so on) are returned as fields of the output structure result. The complete code and data with detailed instructions are provided via a Zenodo archive [29].

**Table 1:** Simulation parameters used in the wingbeat–call synchrony model.

| Parameter | Units | Description |
| --- | --- | --- |
| `bandwidth` | Hz | Two-element vector $[f_{\min}, f_{\max}]$ specifying the frequency range of simulated FM calls. |
| `kr` | dimensionless | Biological reaction constant (responsivity coefficient) relating echo delay to the required call emission interval (Eq. 2) [26]. |
| `target_distance` | m | Initial range to the target at the start of the simulation. |
| `initial_velocity` | m/s | Initial approach velocity of the bat relative to the target. |
| `initial_call_duration` | s | Duration of calls at the beginning of the simulated approach. |
| `max_call_rate` | Hz | Maximum call rate corresponding to empirical buzz call rates. |
| `motile` | logical | Boolean indicating whether the target is stationary or undergoes randomised motion (simulating unpredictable prey). |
| `makeAudio` | logical | Option to generate synthetic audio signals (calls and echoes) in the simulation output. |
| `f_wing` | Hz | Baseline wingbeat frequency used when frequency is fixed. |
| `theta` | rad | Maximum wing excursion angle. |
| `min_theta` | rad | Minimum physiological wing excursion used in the dynamic-θ regime. |
| `max_wingbeat_freq` | Hz | Maximum physiological wingbeat frequency. |
| `wing_sync_mode` | string | Simulation regime: `fixed` (constant wingbeat frequency) or `dynamic` (frequency adapts). |
| `theta_mode` | string | Wing excursion mode: `fixed` (constant excursion) or `dynamic` (excursion adapts to permit higher frequency). |

### 3.4 Parameter Sampling for Comparative Simulations

To further examine how synchrony and asynchrony emerge under different motor-control scenarios, I performed a randomised exploration of biologically plausible parameter combinations. For each of the three simulation conditions (Section 3.3), I generated 200 simulation runs with randomly sampled input parameters and a simulated motile prey that shifted in position with random jitter (*k*_*noise*_ = 0.1). For each sampled simulation run, the following parameters were drawn independently from the distributions specified below: target range (*d*_0_), initial velocity (*v*_0_), responsivity coefficient (*k*_*r*_), baseline wingbeat frequency (*f*_*w*_), and nominal wing excursion angle (*θ*). All remaining parameters were held constant at the values listed in Table 1. Stochasticity therefore entered through the random parameter draws and target jitter, whereas model output was otherwise deterministic for a given sampled parameter set.

The initial target distance was drawn uniformly between 10 and 15 metres, initial flight velocity between 2 and 5 m/s, and responsivity coefficient *k*_*r*_ between 3 and 7. The initial wingbeat frequency was sampled between 3 and 7 Hz, and the maximum excursion angle *θ* was sampled between *π/*6 and *π/*4 radians. To reflect biologically realistic modulation of wingbeat properties, the maximum wingbeat frequency was set as a multiple between 2 and 4 times the baseline frequency, and the minimal excursion amplitude was drawn between 10% and 50% of *θ*. The chosen wingbeat-frequency range spans empirical values reported across seven taxonomic families of microchiroptera [30], with the lower initial values corresponding to larger species in genera such as *Taphozous, Eptesicus* [30], and *Nyctalus* [27]. In each run, all other physiological and acoustic parameters (e.g., sampling frequency, call bandwidth, and call duration) were held constant. This randomised parameter-sampling approach allowed quantification of system behaviour across a broad but biologically relevant parameter space.

### 3.5 Analysis of simulation outputs

Simulation outputs were analysed to characterise how temporal feasibility constraints shape coordination regimes across different motor-control configurations. All analyses were performed on the full ensemble of randomly sampled simulation runs and are descriptive in nature, quantifying distributional structure and effect size.

#### Outcome measures

For each simulation run, the following outcome measures were extracted from the output structure returned by <monospace>simulateEcholocationWings</monospace>:

1. *Number of synchronous calls* (*n*_sync_): the total number of calls classified as synchronous according to the operational definition of synchrony (Section 1.3), requiring accommodation of a single call per wingbeat within the instantaneous excursion limit.
2. *Number of asynchronous calls* (*n*_async_): the complement of *n*_sync_, reflecting calls that could not be accommodated within a wingbeat-aligned emission window.
3. *Fraction of asynchronous calls* (*n*_async_*/n*_total_): a normalised measure that controls for variation in the total number of calls across simulation runs and provides a direct index of coordination breakdown.
4. *Calls per wingbeat*: computed as the ratio between the total number of calls and the total number of wingbeat cycles completed during the simulated approach. This dimensionless quantity provides a continuous measure of departure from one-to-one coordination and captures transitions toward multi-call-per-cycle regimes.
5. *Wingbeat phase of asynchronous calls*: for calls classified as asynchronous, emission times were converted to wingbeat phase fractions using Eq. (7), yielding a circular variable on [0, 1] that characterises the structure of phase slip.

#### Descriptive statistics

For each simulation condition, outcome distributions were summarised using robust descriptive statistics, including medians, interquartile ranges (IQR), 5–95% percentile ranges, and median absolute deviation (MAD). These summaries characterise the typical behaviour of the system and its variability across parameter space without assuming underlying sampling noise or independent biological replicates.

#### Effect size estimation

To quantify the magnitude of distributional shifts between simulation conditions, pairwise comparisons were performed using two complementary effect-size measures:

1. The difference in medians between conditions, with 95% bootstrap confidence intervals obtained by resampling simulation runs with replacement.
2. Cliff’s delta (*δ*), a non-parametric effect size that quantifies the probability that an outcome drawn from one condition exceeds an outcome drawn from another.

These effect sizes provide a scale-free assessment of how strongly coordination regimes differ under alternative motor-control constraints without relying on null-hypothesis significance testing [31]. Because these simulation runs represent deterministic outcomes for sampled parameter combinations rather than independent biological replicates, classical significance tests based on sampling variability are not appropriate. Instead, the goal of the analysis is to quantify the magnitude and direction of distributional differences across model conditions. Cliff’s delta is well suited for this purpose because it is a non-parametric, distribution-free measure that is robust to non-normal and heteroscedastic data and directly expresses the probability that values drawn from one condition exceed those from another. Together with bootstrap confidence intervals on median differences, this approach provides an interpretable description of regime shifts across parameter space without invoking inferential tests that assume stochastic sampling.

Complete numerical summaries and effect-size estimates for all outcome measures are provided in Supplementary Tables A1-A6.

#### 3.5.1 Quantifying phase locking in simulations

To quantify the presence and strength of wingbeat–call phase locking in simulated data, call emission times were converted to wingbeat phases using Eq. (7). The resulting phase angles *{ϕ*_*n*_*}* define a circular variable on [0, 2*π*), allowing standard circular statistics to be applied.

Phase concentration was assessed using the Rayleigh statistic, which tests the null hypothesis that phases are uniformly distributed around the cycle. The mean resultant vector length,

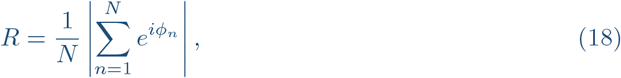

provides a measure of phase locking strength, with *R* ≈ 0 indicating uniform phase dispersion and *R* → 1 indicating strong clustering. Statistical significance was evaluated using the Rayleigh test for non-uniformity.

## 4. RESULTS

### 4.1 Illustrative coordination regimes under bounded motor dynamics

To provide intuition for how constraints on temporal feasibility shape coordination between echolocation call timing and wingbeat dynamics, I first consider representative simulation trajectories under different motor-control configurations. These examples are intended to illustrate the qualitative coordination regimes implied by the theoretical framework. The prevalence and robustness of these regimes are quantified in the following section.

Figure 2 shows time-domain representations of simulated wingbeat motion together with the timing of call emissions for three motor-control configurations: (i) fixed wingbeat frequency and fixed excursion, (ii) dynamic wingbeat frequency with fixed excursion, and (iii) dynamic wingbeat frequency with dynamic excursion. In all cases, call rate increases over time as the simulated bat approaches the target, leading to a progressive reduction in acoustic delay and, consequently, in the minimum feasible interpulse interval.

**Figure 2:**
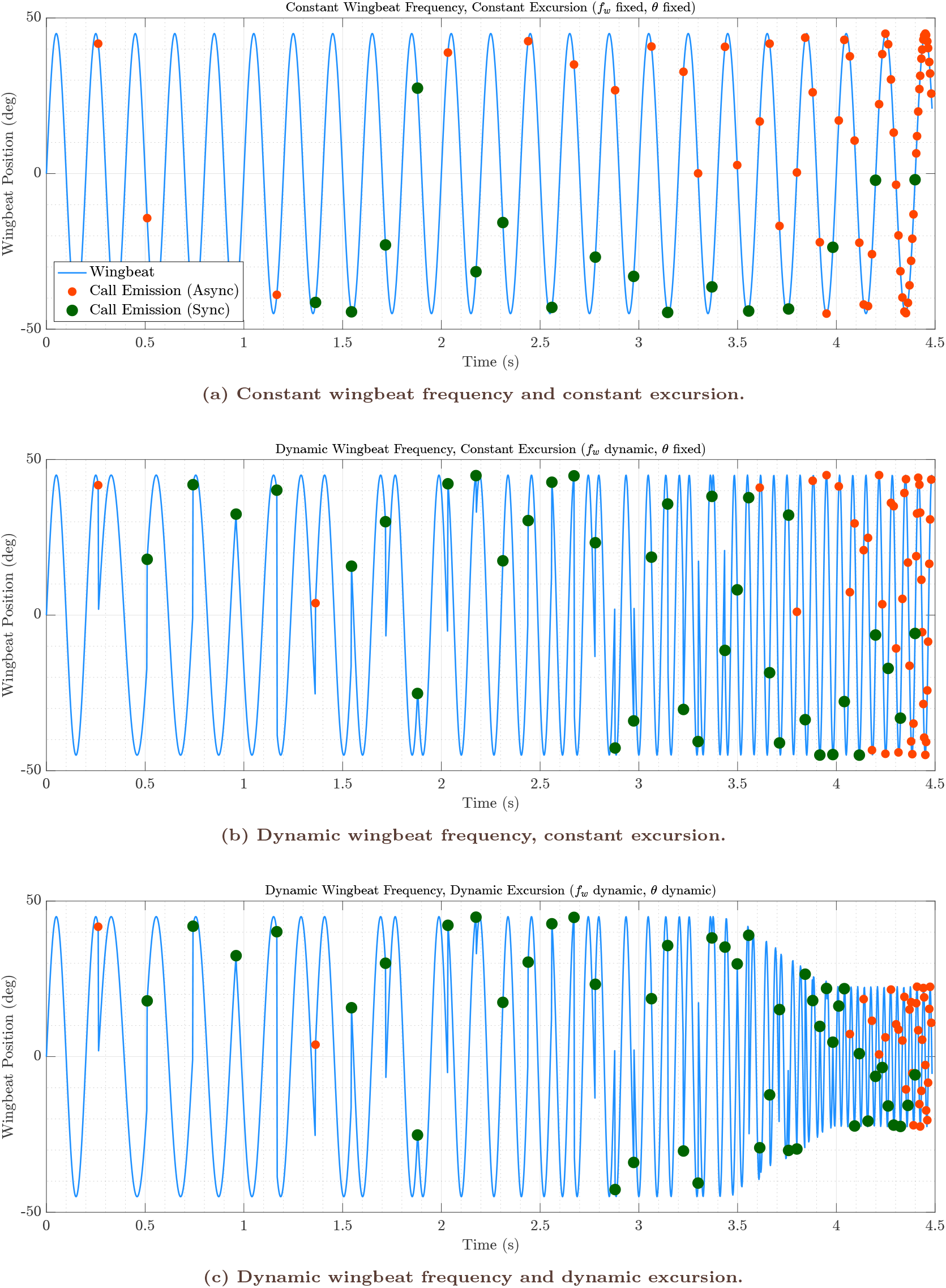
Waveform representations of wingbeat–call synchrony and asynchrony under different motor-control strategies. Each panel shows the simulated wingbeat waveform as a function of time, with circular markers indicating the timing of call emissions. **(A)** With a fixed wingbeat frequency, synchrony between call emission and specific wingbeat phases is maintained only at low call rates; as call rate increases, calls occur at increasingly variable phases. **(B)** When wingbeat frequency is allowed to increase dynamically (up to a maximum), synchrony persists for longer as the bat accelerates its wingbeats to match call timing, but ultimately breaks down as the maximum is reached. **(C)** When both wingbeat frequency and excursion are allowed to adjust, synchrony can be extended, but only within physiological bounds; as call rate surpasses the system’s capacity, asynchrony becomes unavoidable and call emission is dissociated from wing movement. These example trajectories illustrate how precise temporal coordination between call emission and wingbeat is limited by bounded motor dynamics. As sensory demand increases, phase-locked coordination becomes infeasible once physiological limits are reached, giving rise to asynchronous call timing.

Under fixed wingbeat frequency and excursion (Fig. 2a), call emission initially remains phase-locked to a narrow region of the wingbeat cycle at low call rates, but becomes increasingly variable once the wingbeat period can no longer accommodate successive calls within a single cycle. Allowing wingbeat frequency to vary dynamically (Fig. 2b) extends the range of call rates over which phase-locked coordination can be maintained by accelerating the motor rhythm to match the shrinking interpulse interval. Allowing both wingbeat frequency and excursion to vary (Fig. 2c) extends synchrony furthest by enlarging the phase window available for call insertion. Even in this most flexible configuration, however, synchrony persists only while the motor parameters remain within imposed bounds. Across all three conditions, increased motor flexibility shifts the boundary of temporal feasibility but does not eliminate it.

The same pattern is visible in the complementary phase-based representation in Figure 3. There, call timing is expressed as the predicted wingbeat phase at which each call would be inserted on the basis of the current acoustic delay and response window, together with the actual wingbeat phase at call emission. For each motor-control configuration, synchronous calls occupy a bounded region of phase space defined by the available wingbeat excursion. As call rate increases, the predicted insertion phase moves progressively towards this boundary. Once the predicted phase exceeds the admissible range, synchronous insertion is no longer possible and calls occur outside the synchrony-permissive window, producing phase slip and asynchronous timing.

**Figure 3:**
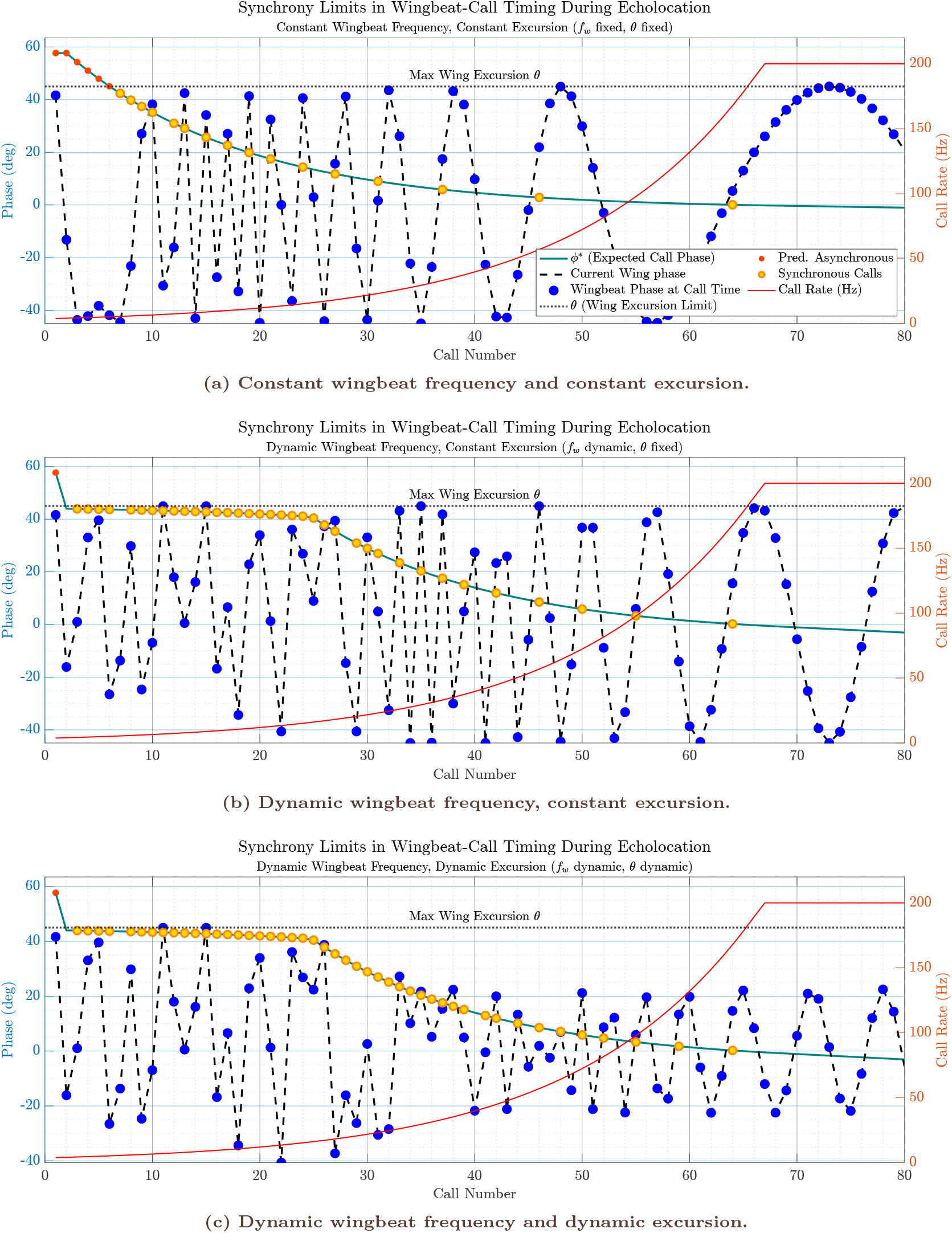
Simulation of wingbeat–call synchrony under increasing sensory-motor demand. Each panel shows the evolution of predicted and actual wingbeat–call phase relationships as a simulated bat approaches a stationary target, with call rate and wing kinematics computed from the theoretical framework (see text). **Teal line**: Predicted call insertion phase (*ϕ*^*^) in degrees, representing the optimal phase within the wingbeat cycle for emitting each call, given the current echo delay and response window. **Blue dots on black dashed line**: Actual phase of the wingbeat at the time of each call emission. **Red dots**: Calls that are asynchronous by the phase-only criterion (predicted phase exceeds the current excursion). **Yellow dots**: Calls that are truly synchronous, i.e., occur within a new wingbeat, at a phase within the current excursion, and at no more than one call per wingbeat. **Red line (right axis)**: Call rate profile (Hz), increasing as the bat nears the target. **Horizontal dashed line**: Maximum (or current) wing excursion (*θ*), serving as the phase limit for synchronous call insertion. (**a**) With fixed wingbeat frequency and fixed excursion, synchrony (green) is possible only at low call rates. As call rate rises, synchrony rapidly breaks down, as wingbeat cannot keep pace with the sensory-motor demand. (**b**) With dynamic wingbeat frequency (but fixed excursion), synchrony persists further into the approach, but is ultimately lost when the physiological limit of wingbeat frequency is reached. (**c**) When both wingbeat frequency and excursion are allowed to adapt (i.e., *θ* decreases as *f*_w_is capped), synchrony can be maintained the longest. Only when both the frequency and amplitude limits are reached does synchrony become impossible, resulting in a rapid onset of asynchrony. These results illustrate the progressive trade-offs bats may employ to maintain temporal coordination of call emission and wingbeat, and reveal how phase-locked coordination breaks down once temporal feasibility bounds are exceeded in high call-rate regimes.

Across all configurations, these illustrative trajectories show that transitions between synchronous and asynchronous coordination arise from the interaction between shrinking sensory delays and bounded motor dynamics. In the next section, I quantify how often these regimes occur across parameter space and how motor flexibility alters the relative prevalence of synchronous and asynchronous coordination.

### 4.2 Distribution of coordination regimes across parameter space

I next quantified how the coordination regimes illustrated in Section 4.1 are expressed across parameter space using ensembles of randomly sampled simulation runs under each motor-control configuration (Section 3.5). Because each simulation run represents a deterministic outcome of specified temporal and motor constraints under parameter variation, the resulting distributions reflect heterogeneity across parameter space rather than stochastic measurement error. Accordingly, I summarise these distributions using robust statistics and distribution-level effect sizes (median differences with bootstrap confidence intervals; Cliff’s delta) to characterise how motor constraints shape the prevalence of synchronous and asynchronous coordination.

#### Motor flexibility reduces asynchronous timing and expands the synchrony-permissive regime

Across simulation runs, the fraction of asynchronous calls per run decreased systematically as motor flexibility increased (Fig. 4a). Under fixed wingbeat frequency and fixed excursion (Condition 1), the model produced predominantly asynchronous timing, with a median asynchronous fraction of 0.8269 (IQR 0.0266) (see Table 2). Allowing wingbeat frequency to vary dynamically while keeping excursion fixed (Condition 2) substantially reduced this proportion (median 0.5714, IQR 0.0969), and allowing both dynamic wingbeat frequency and dynamic excursion (Condition 3) reduced it further (median 0.4600, IQR 0.0833) (also see Supplementary Tables A1-A4). These shifts indicate that increasing motor flexibility enlarges the region of parameter space in which phase-locked call insertion remains temporally feasible, consistent with the qualitative regime progression illustrated in Figs. 2–3.

**Table 2:** Headline coordination metrics across simulation conditions. Values are reported as median (IQR) across *N* = 200 sampled simulation runs per condition. The fraction of asynchronous calls and calls-per-wingbeat provide normalised indicators of regime structure that are robust to differences in total call output across runs. Full descriptive statistics underlying these headline metrics are provided in Supplementary Tables A1-A4.

| Condition | $f_w$ | $\theta$ | Fraction async | Calls per wingbeat |
| --- | --- | --- | --- | --- |
| C1 | fixed | fixed | 0.827 (0.027) | 2.710 (1.418) |
| C2 | dynamic | fixed | 0.571 (0.097) | 1.967 (0.444) |
| C3 | dynamic | dynamic | 0.460 (0.083) | 1.562 (0.213) |

**Figure 4:**
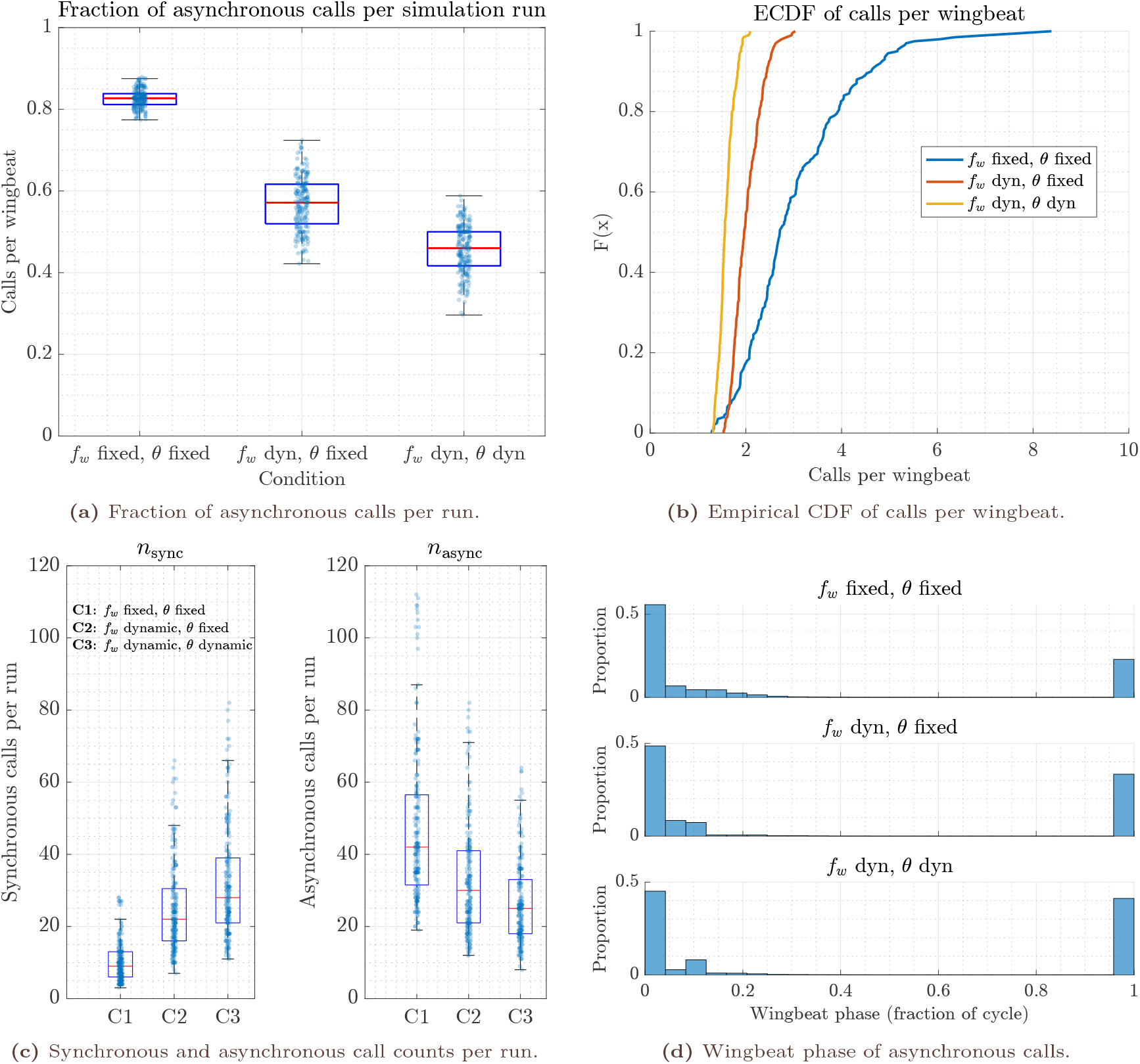
Distributional structure of wingbeat–call coordination regimes across motor-control configurations. (a) Fraction of asynchronous calls per simulation run (median, IQR, whiskers; points denote individual sampled runs). (b) Empirical cumulative distribution functions (ECDFs) of calls per wingbeat, a dimensionless indicator of departure from one-to-one coordination. (c) Raw counts of synchronous (*n*_sync_) and asynchronous (*n*_async_) calls per run, shown for completeness but sensitive to variation in total call output. (d) Wingbeat phase distributions of asynchronous calls (mapped to [0, 1]), pooled across runs within each condition. Across panels, increasing motor flexibility (from fixed wingbeat frequency and excursion to dynamic frequency and excursion) systematically reduces the prevalence of asynchronous timing and shifts the system toward coordination regimes compatible with phase-locked call insertion. Asynchronous calls are not uniformly distributed in phase but cluster near the boundary of the synchrony-permissive window, consistent with structured phase slip arising from temporal feasibility constraints rather than phase-random decoupling. Normalised measures (fraction asynchronous; calls per wingbeat) provide the most robust indicators of regime structure across parameter space.

Pairwise distribution comparisons confirm that these changes are large and robust (median differences with bootstrap CIs and Cliff’s *δ*; see Supplementary Table A5). Relative to Condition 1, Condition 2 reduced the median asynchronous fraction by 0.2555 (95% bootstrap CI [−0.2684, −0.2428]), and Condition 3 reduced it by 0.3669 (95% bootstrap CI [−0.3821, −0.3549]). The corresponding Cliff’s deltas for Condition 1 versus Conditions 2 and 3 were −1, indicating near-complete separation of the outcome distributions under the sampled parameter ensemble. Condition 3 additionally reduced the asynchronous fraction relative to Condition 2 by 0.1114 (95% bootstrap CI [−0.1305, −0.0946]; *δ* = −0.7857), indicating that excursion flexibility provides further, though smaller, expansion of feasible coordination once frequency flexibility is already available.

#### Calls-per-wingbeat decreases with motor flexibility, indicating reduced reliance on multi-call regimes

A complementary indicator of coordination breakdown is the number of calls emitted per wingbeat cycle. Across simulations, calls-per-wingbeat decreased monotonically as motor flexibility increased (Fig. 4b), reflecting reduced reliance on multi-call-per-cycle regimes when wingbeat dynamics can adapt to shrinking temporal feasibility windows (see Supplementary Table A4). Under Condition 1, calls-per-wingbeat were highly variable and frequently exceeded two (median 2.710, IQR 1.418; 5–95% range [1.603, 5.172]). Under Condition 2, the distribution shifted toward lower values (median 1.967, IQR 0.444), and under Condition 3 it shifted lower still (median 1.562, IQR 0.213).

Median differences in calls per wingbeat were −0.743 for Condition 2 relative to Condition 1 (95% CI [−0.944, −0.620]; *δ* = −0.625), −1.148 for Condition 3 relative to Condition 1 (95% CI [−1.347, −1.042]; *δ* = −0.886), and −0.404 for Condition 3 relative to Condition 2 (95% CI [−0.477, −0.340]; *δ* = −0.829) (see Supplementary Table A5). Together, these results show that allowing frequency and excursion adaptation shifts the system toward coordination patterns closer to one-to-one emission within the wingbeat cycle, without implying that such synchrony constitutes a behavioural objective.

### Synchronous and asynchronous call counts shift in opposite directions, but normalised measures are more interpretable

Raw counts of synchronous and asynchronous calls per run varied systematically across motor-control configurations (Fig. 4c). The median number of synchronous calls increased from 9 in Condition 1 (IQR 7) to 22 in Condition 2 (IQR 14.5) and 28 in Condition 3 (IQR 18). Conversely, the median number of asynchronous calls decreased from 42 (IQR 25) to 30 (IQR 20) and 25 (IQR 15), respectively. Because the total number of calls per run can vary across simulations, normalised measures such as the fraction of asynchronous calls and calls-per-wingbeat provide the most directly interpretable indicators of coordination regime structure. Raw event counts are nevertheless retained alongside normalised fractions because they preserve information about the absolute magnitude of call output during each simulated approach. Whereas fractions characterise the relative balance between synchronous and asynchronous regimes, raw counts indicate the total sensory sampling effort generated by the system and therefore provide context for interpreting regime shifts across motor-control configurations.

### Asynchronous call phases exhibit structured phase-slip near feasibility boundaries

To assess whether asynchronous calls in the model occur at arbitrary wingbeat phases or exhibit structured timing, I analysed the wingbeat phase fractions of asynchronous call emissions (Fig. 4d). Across all conditions, asynchronous phases were strongly clustered within a narrow phase interval (mean phase fractions ≈ 0.017–0.030), with high mean resultant lengths (*R* = 0.9367–0.9678). This concentration indicates that asynchrony in the model is dominated by phase-slip events occurring near the boundary of the synchrony-permissive window, rather than by uniformly distributed decoupled timing. This phase structure arises from the boundary-defined synchrony criterion used in the model and should not be interpreted as evidence for preferred biomechanical or respiratory phases in bats. Circular summary statistics for asynchronous call phases are reported in Supplementary Table A6.

## 5 DISCUSSION

### 5.1 Temporal feasibility as a governing constraint on closed-loop echolocation

Bat echolocation is a canonical closed-loop sensing system: vocal emissions shape the sensory scene, echoes return after a propagation delay, and the animal updates behaviour under finite neural and motor latencies [3, 5, 8, 32]. The present study formalises a narrower question: *when is closed-loop timing compatible with bounded response dynamics and an ongoing cyclic motor rhythm?* In the framework developed here, that question is answered by feasibility relations that specify the shortest admissible interpulse interval (IPI) consistent with echo-guided updating (Eq. 4 and related expressions), and by the consequent compatibility conditions for phase-locked coordination with a wingbeat rhythm (e.g. Eq. 9 and the phase-violation criterion in Eq. 14).

A key conceptual point is that feasibility is not a behavioural objective. The model does not assume that bats *seek* wingbeat–call synchrony, minimise asynchrony, or enforce a particular phase. Rather, the framework delineates a region of state space in which phase-locked insertion is possible and a region in which it is not, independent of motivation or strategy. In this sense, the work provides a constraint-based baseline: it separates patterns that could plausibly be produced by any controller from patterns that would require additional mechanisms (or would be impossible) once temporal margins become binding.

### 5.2 Buffered pseudo–closed-loop behaviour can masquerade as feedback control

A call–echo–call sequence is not, by itself, sufficient evidence for tight closed-loop control. When acoustic delays are short relative to the animal’s effective update time, echoes often arrive with substantial temporal slack. In this buffered regime, a bat could emit calls according to an internally generated schedule or a kinematic heuristic, while still receiving informative echoes “in time” before the next emission becomes mechanically or neurally available. Such *pseudo– closed-loop* operation can therefore mimic echo-anchored regulation without requiring that call timing be causally governed by echo arrival. This interpretation is consistent with recent analyses emphasising that, in some contexts, call timing can be accurately generated by internal pacing mechanisms when echoes are not rate-limiting [33].

The distinction becomes behaviourally consequential once the buffer collapses. As ranges shorten and call rates rise, the informative portion of the echo response competes directly with the scheduling of subsequent emissions. In the responsivity formulation, this corresponds to the regime in which IPI is explicitly constrained by the combination of echo delay and the bounded response window (Eq. 4–6). Importantly, this does not imply that bats must wait for the full termination of the echo stream: the relevant anchor is the earliest time at which sufficient *information* is available to drive an update. Nevertheless, once the feasibility bound is active, internally paced schedules can no longer remain decoupled from the acoustic scene.

Within the buffered regime, apparent invariants such as an approximately constant *distance travelled per call* can arise as emergent regularities rather than controlled objectives. If call timing follows an internal policy IPI ≈ *g*(*v*) that covaries with velocity over a restricted behavioural range, the derived spatial sampling interval — the product of instantaneous velocity and interpulse interval — can appear stable even when echo timing is not rate-limiting. This provides a concrete route by which “invariants” can be observed under conditions where propagation delays are not yet the dominant constraint.

In contrast, once the system enters the temporally constrained regime (Eq. 5 and the saturation bound in Eq. 6), maintaining a fixed spatial sampling interval becomes progressively incompatible with the necessary contraction of IPI with decreasing range. The prediction is therefore not that invariants are “excluded”, but that they should be *regime contingent*: they are most expected under buffered operation and should degrade systematically as temporal feasibility becomes the binding limitation, particularly in high-demand phases such as pursuit and the terminal buzz, as the results from Jakobsen et. al. (2025) indicate [25]. Moreover, when temporal overlap threatens interpretability (e.g. increasing broadcast–echo ambiguity in clutter), bats are known to recruit signal-domain degrees of freedom rather than attempting to preserve a fixed spatial sampling interval; one concrete example is dynamic frequency shifting to reduce ambiguity under cluttered echo streams [34]. In the present framing, such adjustments are naturally interpreted as auxiliary control dimensions recruited when feasibility constraints begin to bind, rather than as evidence that a single spatial invariant is the maintained objective across regimes.

### 5.3 Synchrony and asynchrony as regime outcomes rather than control objectives

The framework provides a parsimonious interpretation of wingbeat–call synchrony. In low-demand conditions (longer delays and longer feasible IPIs), there is ample slack: many call insertion times are compatible with closed-loop updating, and one-to-one locking can occur without requiring a dedicated coupling mechanism. Synchrony is therefore expected during steady flight and other low-demand tasks, consistent with long-standing observations of phaselocked calling in some contexts [20–22]. At broader comparative scales, scaling relationships between wingbeat frequency and pulse emission rate, including common 1:1 and 1:2 coordination patterns, likewise support the view that phase-locked regimes are feasible and often realised under permissive conditions, without implying that phase locking itself is the control variable [35].

As sensory demand increases, however, feasibility margins contract. The wingbeat period cannot be reduced indefinitely, and the admissible phase window for inserting calls within a wing excursion narrows (Eq. 12–15). Under these conditions, strict phase locking becomes fragile and eventually infeasible. Crucially, this transition does not require a discrete switch in behavioural mode. Rather, coupling is expected to degrade progressively as call rate increases and the temporal margins available for motor coordination collapse. This interpretation aligns with empirical reports that coupling is common at low duty cycles but degrades as call rate increases, and that wild bats can transiently decouple sound production from wingbeats during demanding capture manoeuvres [23, 24].

The term “brief decoupling” as used by Stidsholt et al. (2021) [23] can therefore be interpreted precisely within the present framework. In their data, decoupling does not correspond to a sustained loss of coordination, but to short intervals—predominantly during late approach and the buzz—in which call timing no longer exhibits a stable phase relationship to the wingbeat cycle.

The feasibility analysis developed here predicts exactly such transient breakdowns. The simulation model, built on the core identities of the responsivity framework [26], reproduces this pattern while keeping call timing strictly echo-constrained. In that sense, the observed decoupling is more parsimoniously interpreted as a consequence of closed-loop timing than as an active disengagement of wingbeat–call coordination.

Recent laboratory evidence further supports this interpretation. Xia et al. (2025) show that wingbeat–call coupling is flexible across species, call types, and task demands, with structured synchrony during permissive regimes and systematic decoupling during high call-rate sequences such as long sonar sound groups and terminal buzzes [24]. Taken together, the field observations of Stidsholt et al. and the controlled laboratory results of Xia et al. suggest that wingbeat– call decoupling need not represent an optional behavioural strategy. Instead, it arises naturally when temporal constraints imposed by closed-loop sensing exceed what a cyclic motor rhythm can accommodate.

This interpretation is especially relevant during late approach and interception, when the feasible interpulse interval contracts rapidly while flight mechanics are also being reconfigured for prey capture [5, 36–39]. For a stationary reflector, *T*_*a*_ provides a reliable estimate of range, and successive emissions can be planned from information that remains valid across the IPI. For an erratically moving prey item, by contrast, the echo-derived state available at emission time is already lagged by the time the next call is scheduled. Maintaining strict wingbeat–call synchrony under those conditions would require continuous compensatory adjustments of flight kinematics precisely when the feasibility window is shrinking. The disappearance of synchrony during pursuit therefore need not imply a loss of control; it may simply indicate that phase locking has become infeasible given the prevailing delays, response bounds, and motor limits.

The simulations further predict that asynchrony need not be phase-random. When the feasibility boundary is defined relative to a limited excursion window, violations occur preferentially near that boundary, producing structured phase slip. In this study, the clustering of asynchronous phases is therefore interpreted as a *geometric consequence of the feasibility criterion* (Eq. 14–15), not as evidence for preferred biomechanical or respiratory phases in bats. This distinction matters because structured asynchrony can arise even within a single underlying control architecture simply because the admissible timing region has collapsed.

### 5.4 Comparative and morphological implications

Although the model is intentionally agnostic about detailed aerodynamics and muscle physiology, it yields clear comparative implications. Morphological and biomechanical differences should primarily shift *where* feasibility boundaries occur, rather than introducing fundamentally different coordination principles. Species with broader kinematic ranges (e.g. higher sustainable wingbeat frequencies, greater flexibility in excursion) should retain phase-locked coordination over a wider range of echo delays and call rates, whereas species with constrained flight dynamics should cross feasibility limits earlier. This perspective is consistent with the wide diversity of flight morphology and foraging guilds in bats, and with the continuous variation in echolocation behaviour across ecological contexts [5, 24, 40, 41]. Comparative scaling of wingbeat frequency and pulse emission rate provides an additional constraint-based bridge between morphology and coordination regimes, supporting the interpretation that morphological differences modulate the location of feasibility boundaries rather than altering the governing principles [35].

Notably, the framework does *not* ascribe adaptive value to synchrony or asynchrony themselves. Instead, it predicts quantitative shifts in the prevalence of coordination regimes under comparable sensory demands. This makes the comparative question tractable: measure how the synchrony-permissive region moves with morphology and kinematic capacity, rather than asking whether a species “uses synchrony” as a strategy.

An additional implication of the simulations is that flexibility in wingbeat frequency expands the synchrony-permissive regime more strongly than flexibility in wing excursion. This difference follows directly from the distinct roles played by these variables in the timing structure of the system. Wingbeat frequency determines the duration of the motor cycle (*T*_*w*_) and therefore sets the temporal capacity of the system to accommodate successive calls. Increasing frequency effectively shortens the cycle, allowing additional opportunities for call insertion before the interpulse interval becomes shorter than the wingbeat period. By contrast, changes in wing excursion modify only the admissible phase window within a given cycle. Although larger excursion limits can slightly extend the range of phases in which synchronous calls can occur, they do not alter the fundamental time-bounds imposed by the wingbeat period itself. This asymmetry highlights that the primary control variable governing coordination capacity is the temporal spacing of motor cycles, whereas excursion mainly affects the tolerance of phase alignment within each cycle.

### 5.5 Methodological implications for interpreting coordination data

The feasibility framework clarifies which aspects of wingbeat–call coordination are diagnostically informative and which are not. Because phase-locked calling can occur whenever temporal margins are permissive, the mere presence of synchrony is insufficient evidence for causal motor coupling. Conversely, asynchrony is an expected outcome once feasibility bounds are exceeded and therefore does not, by itself, imply flexible or decoupled control. Interpreting coordination data therefore requires explicit reference to the temporal regime in which calls are produced.

The framework also specifies where additional measurements are essential. Acoustic timing alone cannot distinguish buffered pseudo–closed-loop operation from genuinely echo-limited control. Disambiguating these regimes requires concurrent kinematic data, particularly wingbeat phase and excursion, obtained via high-speed videography, inertial sensors, or on-board telemetry. Recent developments in on-board telemetry provide a powerful means to elucidate the temporal coordination of behavioural parameters in freely flying bats [42].

More generally, the feasibility analysis motivates a shift in emphasis from asking whether calls are synchronised to asking *when synchrony is possible at all*. By anchoring coordination metrics to explicit temporal constraints, empirical studies can avoid over-interpreting correlation structure and instead test whether observed patterns track the boundaries imposed by delayed feedback and bounded response dynamics.

### 5.6 Generality, limitations, and testable predictions

The logic of temporal feasibility is not specific to bats. Any active sensing system that relies on self-generated signals, delayed feedback, and bounded response dynamics will exhibit analogous constraints, including electrosensing fish and whisking rodents [17, 43], as well as engineered active-perception systems [14]. In such systems, coordination with rhythmic actuation is expected to hold in permissive regimes and to fail in structured ways once temporal margins collapse.

The present work deliberately omits energetic costs, detailed aerodynamics, sensory noise, and learning. Accordingly, it should be read as identifying *necessary constraints*, not as a complete behavioural model. Within that scope, it yields testable predictions: (i) synchrony breakdown should correlate more strongly with temporal demand (echo delay, required response window) than with categorical behavioural labels; (ii) feasibility boundaries should shift with kinematic capacity across individuals and species; and (iii) asynchronous timing near transitions should show structured phase-slip consistent with the geometry implied by Eq. 14–15. These predictions can be evaluated with combined acoustic and kinematic datasets collected across controlled range manipulations and natural pursuit sequences.

The present framework is mechanistic rather than inferential: it derives expected timing relations from the closed-loop coupling between call emission, echo return, processing delay, and motor dynamics. By contrast, causality-analysis methods such as Granger causality [44] quantify whether past values of one measured variable improve prediction of another. These approaches are complementary. For example, simultaneous call-timing and wingbeat time series could be analysed to test whether the lagged directional dependencies inferred statistically are consistent with the temporal ordering and delay structure predicted here.

## 6 CONCLUSIONS

This study identifies *temporal feasibility* as a fundamental constraint shaping wingbeat–call coordination in bat echolocation. By formalising the timing dependencies between call emission, acoustic delay, sensory processing, and motor response, the framework specifies when phaselocked coordination with a rhythmic motor cycle is possible and when it must fail. Synchrony and asynchrony thus emerge as regime-dependent outcomes of bounded response dynamics, rather than as behavioural objectives or discrete control strategies.

Under temporally permissive conditions, substantial slack allows phase-locked calling to occur without requiring dedicated coupling mechanisms. As sensory demand increases during pursuit and the terminal buzz, feasibility margins contract and the admissible phase window for call insertion collapses relative to the wingbeat period. The resulting phase slip and brief decoupling arise necessarily, even when call timing remains echo-constrained. This provides a mechanistic explanation for empirical observations of transient decoupling during capture manoeuvres without invoking changes in control architecture.

By separating feasibility constraints from behavioural goals, the framework reconciles diverse empirical patterns, including phase locking, discrete coordination modes, and regime-specific breakdown of synchrony. More broadly, the results illustrate how rhythmic motor coordination in active sensing systems is governed by delayed feedback and bounded response dynamics, offering a general constraint-based perspective for interpreting sensorimotor timing in ecological behaviour.

## AUTHOR CONTRIBUTIONS

The study was conceived, conducted, and written by the sole author.

## ACKNOWLEDGEMENTS

I thank my colleagues at the Technical University of Munich for helpful discussions and support during the development of this work. I also thank the anonymous reviewers for their constructive comments, which helped improve and clarify the manuscript.

## CONFLICTS OF INTEREST

The author declares no competing interests.

## DATA AVAILABILITY STATEMENT

The simulation code, experimental data, and analysis scripts supporting the findings of this study are publicly available at Zenodo DOI: https://doi.org/10.5281/zenodo.15854977

## FUNDING

This study did not use external funding.

### Manuscript Information

Version: R2

Peer-Review 1: September 24, 2025

Peer-Review 2: March 03, 2026

Last Compiled: April 24, 2026

Words in text: 8009

Number of floats/tables/figures: 6

Status: Accepted by the journal *Ecology and Evolution*

## APPENDIX

## A SUPPLEMENTARY TABLES

**Table A1:** Descriptive statistics for synchronous call counts per run. Summary statistics are computed across *N* = 200 sampled simulation runs for each condition.

| Condition | Mean | SD | Median | IQR | P05 | P25 | P75 | P95 | MAD |
| --- | --- | --- | --- | --- | --- | --- | --- | --- | --- |
| C1 ($f_w$ fixed, $\theta$ fixed) | 10.2 | 5.6 | 9 | 7.0 | 4.0 | 6.0 | 13.0 | 21.5 | 3 |
| C2 ($f_w$ dyn, $\theta$ fixed) | 24.9 | 12.3 | 22 | 14.5 | 10.0 | 16.0 | 30.5 | 50.5 | 7 |
| C3 ($f_w$ dyn, $\theta$ dyn) | 32.0 | 15.3 | 28 | 18.0 | 14.0 | 21.0 | 39.0 | 64.0 | 9 |

**Table A2:** Descriptive statistics for asynchronous call counts per run. Summary statistics are computed across *N* = 200 sampled simulation runs for each condition.

| Condition | Mean | SD | Median | IQR | P05 | P25 | P75 | P95 | MAD |
| --- | --- | --- | --- | --- | --- | --- | --- | --- | --- |
| C1 ($f_w$ fixed, $\theta$ fixed) | 46.9 | 21.2 | 42 | 25.0 | 24.0 | 31.5 | 56.5 | 100.5 | 12 |
| C2 ($f_w$ dyn, $\theta$ fixed) | 32.6 | 15.2 | 30 | 20.0 | 14.5 | 21.0 | 41.0 | 67.5 | 10 |
| C3 ($f_w$ dyn, $\theta$ dyn) | 26.5 | 11.6 | 25 | 15.0 | 12.5 | 18.0 | 33.0 | 50.0 | 7 |

**Table A3:** Descriptive statistics for asynchronous call fraction per run. Summary statistics are computed across *N* = 200 sampled simulation runs for each condition.

| Condition | Mean | SD | Median | IQR | P05 | P25 | P75 | P95 | MAD |
| --- | --- | --- | --- | --- | --- | --- | --- | --- | --- |
| C1 ($f_w$ fixed, $\theta$ fixed) | 0.827 | 0.023 | 0.827 | 0.027 | 0.790 | 0.812 | 0.838 | 0.866 | 0.013 |
| C2 ($f_w$ dyn, $\theta$ fixed) | 0.569 | 0.065 | 0.571 | 0.097 | 0.467 | 0.520 | 0.617 | 0.672 | 0.047 |
| C3 ($f_w$ dyn, $\theta$ dyn) | 0.457 | 0.059 | 0.460 | 0.083 | 0.356 | 0.417 | 0.500 | 0.547 | 0.041 |

**Table A4:** Descriptive statistics for calls per wingbeat. Summary statistics are computed across *N* = 200 sampled simulation runs for each condition.

| Condition | Mean | SD | Median | IQR | P05 | P25 | P75 | P95 | MAD |
| --- | --- | --- | --- | --- | --- | --- | --- | --- | --- |
| C1 ($f_w$ fixed, $\theta$ fixed) | 3.028 | 1.178 | 2.710 | 1.418 | 1.603 | 2.214 | 3.633 | 5.172 | 0.662 |
| C2 ($f_w$ dyn, $\theta$ fixed) | 2.024 | 0.305 | 1.967 | 0.444 | 1.622 | 1.783 | 2.227 | 2.556 | 0.214 |
| C3 ($f_w$ dyn, $\theta$ dyn) | 1.587 | 0.166 | 1.562 | 0.213 | 1.344 | 1.475 | 1.688 | 1.898 | 0.102 |

**Table A5:** Pairwise effect sizes between conditions. For each metric, the table reports the difference in medians (CondA−CondB) with bootstrap 95% confidence intervals, and Cliff’s *δ* with 95% confidence intervals.

| Metric | Comparison | ΔMedian | CI<sub>low</sub> | CI<sub>high</sub> | $\delta$ | CI<sub>low</sub> | CI<sub>high</sub> |
| --- | --- | --- | --- | --- | --- | --- | --- |
| Synchronous calls ($n_{\text{sync}}$) | C1–C2 | 13.0 | 11.0 | 15.0 | 0.800 | 0.740 | 0.854 |
| Synchronous calls ($n_{\text{sync}}$) | C1–C3 | 19.0 | 16.0 | 22.0 | 0.908 | 0.872 | 0.941 |
| Synchronous calls ($n_{\text{sync}}$) | C2–C3 | 6.0 | 3.0 | 9.5 | 0.302 | 0.198 | 0.410 |
| Asynchronous calls ($n_{\text{async}}$) | C1–C2 | -12.0 | -16.0 | -8.0 | -0.450 | -0.547 | -0.348 |
| Asynchronous calls ($n_{\text{async}}$) | C1–C3 | -17.0 | -20.0 | -14.0 | -0.661 | -0.733 | -0.581 |
| Asynchronous calls ($n_{\text{async}}$) | C2–C3 | -5.0 | -9.0 | -1.0 | -0.240 | -0.345 | -0.126 |
| Async fraction ($f_{\text{async}}$) | C1–C2 | -0.255 | -0.268 | -0.243 | -1.000 | -1.000 | -1.000 |
| Async fraction ($f_{\text{async}}$) | C1–C3 | -0.367 | -0.382 | -0.355 | -1.000 | -1.000 | -1.000 |
| Async fraction ($f_{\text{async}}$) | C2–C3 | -0.111 | -0.131 | -0.094 | -0.786 | -0.842 | -0.722 |
| Calls per wingbeat ($c_{\text{pw}}$) | C1–C2 | -0.743 | -0.944 | -0.620 | -0.625 | -0.713 | -0.534 |
| Calls per wingbeat ($c_{\text{pw}}$) | C1–C3 | -1.148 | -1.347 | -1.042 | -0.886 | -0.935 | -0.829 |
| Calls per wingbeat ($c_{\text{pw}}$) | C2–C3 | -0.404 | -0.477 | -0.340 | -0.829 | -0.876 | -0.774 |

**Table A6:** Circular summaries of asynchronous call phases. Asynchronous call phases are pooled within each condition (total *N*_async phases_ shown). Reported are the mean angle (radians), mean phase as a fraction of the wingbeat cycle, mean resultant length *R*, and circular standard deviation.

| Condition | $N_{\text{async}}$ | Mean angle (rad) | Mean phase (frac.) | $R$ | Circ. SD (rad) |
| --- | --- | --- | --- | --- | --- |
| C1 ($f_w$ fixed, $\theta$ fixed) | 9377 | 0.188 | 0.0299 | 0.937 | 0.362 |
| C2 ($f_w$ dyn, $\theta$ fixed) | 6516 | 0.130 | 0.0207 | 0.968 | 0.256 |
| C3 ($f_w$ dyn, $\theta$ dyn) | 5292 | 0.108 | 0.0172 | 0.966 | 0.265 |

## Notes

https://doi.org/10.5281/zenodo.15854977

### Competing Interest Statement

The authors have declared no competing interest.

### Summary of Updates

Following the second revision of the manuscript, the article underwent peer review. The subsequent revisions were minor, primarily focusing on presentation, semantics, and grammar. The central thesis of the study remains unchanged from the previous version.

